# SBOL-OWL: An ontological approach for formal and semantic representation of synthetic genetic circuits

**DOI:** 10.1101/499970

**Authors:** Göksel Mısırlı, Renee Taylor, Angel Goñi-Moreno, James Alastair McLaughlin, Chris Myers, John H Gennari, Phillip Lord, Anil Wipat

## Abstract

Standard representation of data is key for the reproducibility of designs in synthetic biology. The Synthetic Biology Open Language (SBOL) has already emerged as a data standard to represent genetic circuit designs, and it is based on capturing data using graphs. The language provides the syntax using a free text document which is accessible to humans only. Here, we provide SBOL-OWL, an ontology for a machine understandable definition of SBOL. This ontology acts as a semantic layer for genetic circuit designs. As a result, computational tools can understand the meaning of design entities in addition to parsing structured SBOL data. SBOL-OWL not only describes how genetic circuits can be constructed computationally, it also facilitates the use of several existing Semantic Web tooling for synthetic biology. Here, we demonstrate some of these features, for example, to validate designs and check for inconsistencies. Through the use of SBOL-OWL, queries are simplified and become more intuitive. Moreover, existing reasoners can be used to infer information about genetic circuit designs that can’t be directly retrieved using existing querying mechanisms. This ontological representation of the SBOL standard provides a new perspective to the verification, representation and querying of information about synthetic genetic circuits and is important to incorporate complex design information via the integration of biological ontologies.

## 1 Introduction

Synthetic biology deals with the rational design, and implementation, of novel biological functions in living systems (*1*). Applications of such engineered systems often rely on synthetic regulatory networks, the so-called genetic circuits, that include different biological parts and complex relationships between them. Reproducibility of these circuits is often challenging due to not having complete information of the designs (*2*). Moreover, design information is often captured using free text, which aids understanding for humans but may be open to interpretation. It is crucial that designs are unambiguously represented and sufficient information is provided for the sake of reproducibility (*3, 4*). This process requires not only capturing data using a common syntax but also agreeing on semantics for machine interoperability. The use of computationally tractable representations of designs, for instance via the GenBank format (*5*), often omit essential information about the descriptions of gene products, such as molecular interactions and hierarchical structure. Visual representations are useful for communicating design function, but are oversimplified and do not reveal detailed information. As the size and complexity of designs increase, it becomes even more important to use automated approaches and to provide machine accessible data.

The Synthetic Biology Open Language (SBOL) (*6, 7*) has emerged as a data format to electronically exchange information about genetic circuits. By using SBOL, data about biological roles, structures and molecular interactions of biological parts such as proteins, DNA, RNA, small molecules, and complex molecules can be captured. Genetic circuits can be represented hierarchically using part-subpart relationships. That is, a parent part can include several child components, which in turn can be composed of other parts. A similar representation is also used to capture molecular interactions in the form of hierarchical modules. This representation is a graph structure with repeated relationships between parent and child components. Graphs are formed of nodes and edges. Nodes can represent entities in a specific domain and edges represent the relationships between these nodes. This approach is ideal to capture complex design information. As a result, the SBOL community adopted a graph-based representation of data, in the form of Resource Description Framework (RDF) documents.

RDF documents are simply graphs, in which domain specific entities can be globally identified using Uniform Resource Identifiers (URIs) and can have properties whose values are typically literals such as strings and integers. Values can also be URIs pointing to other entities. As a result, RDF graphs are defined with triples that include information about an entity of interest, a value, and how this entity and the value is associated.

Using the RDF approach allows SBOL to be flexible when capturing different types of design information in the form of nodes and edges. However, RDF only defines the syntax to represent design information. Tools are still left with the big task of interpreting this information. As the amount of data produced by different research groups in different geographical locations increases, it is becoming more important to capture properties linking different entities and values semantically to make the design information machine understandable. Adopting a graph representation of data also brings the advantage of being able to incorporate custom metadata or annotations. These annotations can either be embedded within SBOL entities to provide additional information or can be used to describe custom domain specific entities. However, the semantics of such metadata are not captured by the SBOL specification. It is therefore important to semantically identify non-SBOL entities and to facilitate their formal representation where possible.

Once a genetic circuit design is constructed, it is crucial to verify the design by querying various design constraints. Graph representation is ideal for SPARQL^1^ (*8*) queries, which are themselves are graph patterns and report matching graph data. However, these queries are created using the RDF syntax and specify exact relationships between nodes and edges. On the other hand logical queries can facilitate writing simpler and intuitive queries that can be used instead of graph-based queries to integrate querying information over multiple types of nodes and edges. Moreover, graph-based queries may not be ideal where hierarchical data are represented through complex relationships, such as is often the case for SBOL.

There have been debates about the particular format of the serialization for SBOL documents. A fully semantic version of SBOL was considered (*9*). As a graph language, RDF has different formats, one of which is eXtensible Markup Language (XML). The RDF/XML format has the advantage of representing graphs for XML tools. As a result, the SBOL community agreed on the RDF/XML serialization to utilize both RDF and XML tools. The rules defining this serialization process is defined in SBOL specifications. These documents are produced as a community effort led by the SBOL chair and editors, and the steering committee. As of this writing, the specification is 132 pages (version 2.2.1), and grows incrementally between different versions. Currently, SBOL data are created using bespoke software libraries and there is no formal representation of the rules to construct SBOL compliant descriptions of genetic circuit designs. Although the RDF/XML format facilitate the graph-based representation of genetic circuit designs, whilst supporting XML tools, providing formal definitions of SBOL entities and their relationships remains an issue.

Currently, rules to validate SBOL documents are written in a free-text specification. These rules are interpreted by software developers, and programmatic validation strategies are applied. Libraries have already been developed in Java, C, Python and JavaScript programming languages. For example, libSBOLj (*10*) is a Java API for reading and writing SBOL documents, and for manipulating SBOL entities. However, the use of such bespoke software libraries is not always necessary to work with SBOL information. As RDF graphs, genetic design information captured using the SBOL data model is already exposed to some of the existing Semantic Web (*11*) resources. Triplestores are used to store design information, and provenance of designs are captured using previously developed Semantic Web resources. The SynBioHub (*12, 13*) design repository, for example, is based on an RDF triplestore, and, hence, standard SPARQL (*14*) graph queries are used to extract design information.

There is clearly a need to semantically enrich the SBOL specification for machine access in order to utilize different tools. Moreover, this process should be flexible to facilitate the development of extensions and to formally capture additional biological design information. Ontologies can provide computable semantics, which can be used by tools to infer new information about design components and their relationships. Gruber defines an ontology as “*an explicit and formal specification of a conceptualisation*”(*15*). An ontological representation of the SBOL data model can also be used to provide a shared understanding of design information in order to capture different entities consistently and unambiguously for machine access.

Ontologies have already been widely used to model biological knowledge and to infer information (*16–19*). However, SBOL entities and their relationships, including those with external terms, are not formally defined in other ontologies. SBOL currently utilizes external ontological terms to standardize the meaning of some of the design information. The SBOL specification (*7, 20*) recommends the use of a set of terms from different ontologies, and lists these terms in human readable tables. These terms often act as values for SBOL specific entities. For example, the `type` property of a DNA component in SBOL should come from the BioPAX ontology (*21*). It is desirable to provide formal definitions of SBOL entities and their relationships ontologically in order to bridge the use of different biological ontologies and genetic design information.

Here, we present SBOL-OWL, an ontological representation of the SBOL specification for machine access. Our goal is to bring a new perspective to the verification, representation and querying of information about synthetic genetic circuits using this ontological approach. The development of libraries has been a key in the adoption of the standard. SBOL-OWL brings another opportunity of using readily available ontological tools such as automated reasoners and ontology editors (*22–24*). This semantic representation of SBOL data also allows richer queries to be expressed in a simpler and more logical manner. Domain specific information can hence be represented as logical axioms that are formed of multiple graph nodes and edges. With this approach, users can query the data more intuitively, rather than with technologies like SPARQL that use complex graph structures to extract information. Moreover, SBOL-OWL facilitates writing ontological queries that cannot be supported using a graph-based approach. In Section 3, we provide examples of such queries, particularly to create recursive queries to fetch information from hierarchical designs. SBOL-OWL facilitates formalizing the SBOL specification for machine interoperability. Such machine access can be used to represent designs semantically, find inconsistencies, execute richer queries, visualize design information using semantic graphs and track changes between different versions of the standard.

## 2 SBOL-OWL: The Synthetic Biology Open Language Ontology

The SBOL-OWL ontology has been developed to provide machine accessible description of the SBOL data standard. The ontology is encoded using the Web Ontology Language (OWL)^2^(*25*), and comes with standard terms or classes. Classes are basic units of ontologies (*16*) and are used to define types of objects in a domain. Here, they represent SBOL specific entities. SBOL-OWL does not change how SBOL is currently used. Instead, it provides a semantically-aware layer for genetic circuit designs. As a result, Semantic Web tools that process OWL files can now be used with genetic circuit designs. For example, OWL reasoners can infer implicit information about these designs using logical axioms. Similarly, OWL tools can better visualize implicit relationships between SBOL-OWL entities.

In the Semantic Web stack, ontologies are used to provide semantics of domain entities whilst RDF is ideal to syntactically represent information. Here, SBOL specific terms form a basis to formalize the standard in the form of the SBOL-OWL ontology and RDF is used to exchange genetic circuit designs. This semantic layer allows tools to understand complex relationships between individual design components and to carry out subsequent processing, such as querying and integrating data.

### 2.1 Encoding SBOL entities

SBOL-OWL terms were created using the free text information in the SBOL data standard. Each term has a unique identifier which corresponds to how an SBOL entity is serialized. Free text meanings of these SBOL entities in the SBOL specification are used to populate terms’ descriptions. The standard `rdfs:label` and `rdfs:comment` properties are used to represent names and descriptions of these terms respectively.

The most important SBOL entities are categorized as `TopLevel` entities, which include `ComponentDefinition`, `ModuleDefinition`, `Sequence`, `Model`, `Attachment`, `Implementation` and `CombinatorialDerivation`. As shown in Figure 1, these `TopLevel` entities are regarded as core terms in the ontology, since these entities are used as containers to exchange information using SBOL.

**Figure 1:**
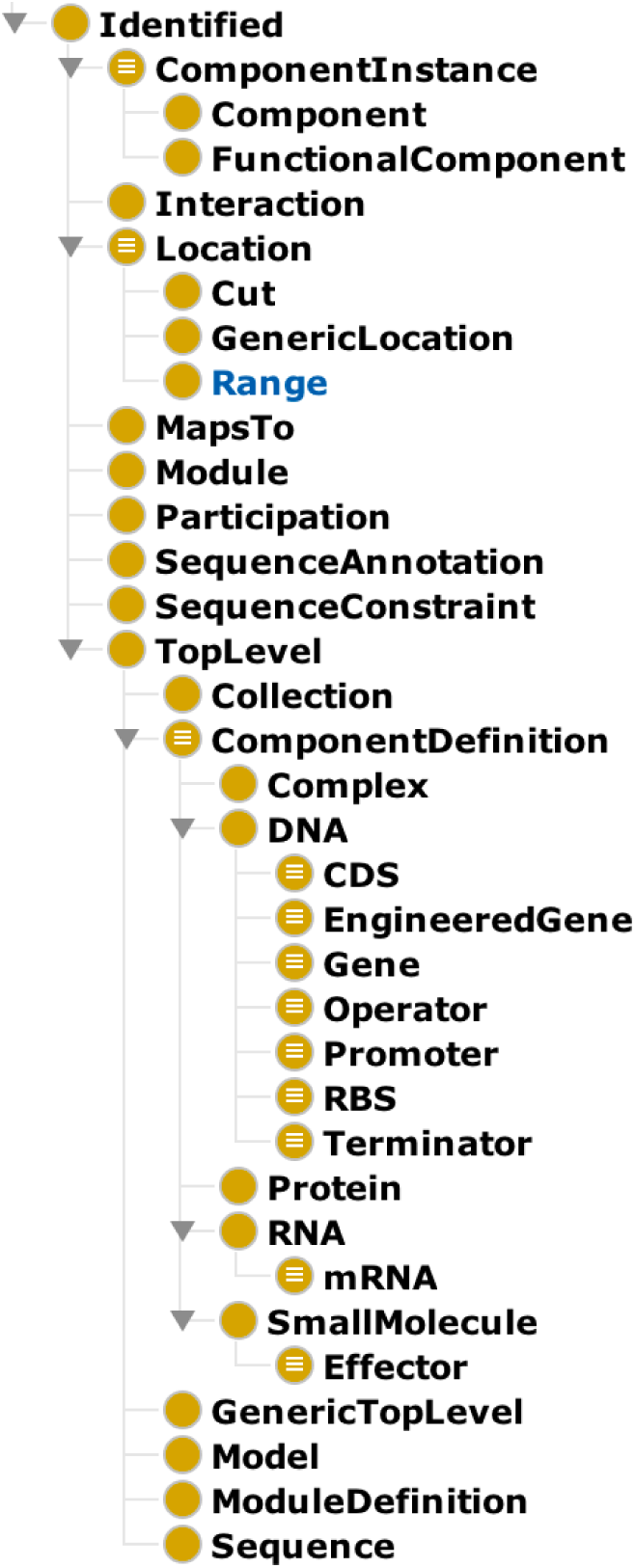
An overview of some of the classes in the SBOL-OWL ontology. Terms are created using SBOL entities that are represented in SBOL documents.

SBOL is a flexible language to represent relationships between different design compo nents and modules in order to represent complex and hierarchical designs. There are several entities that are used to provide additional important information and are also represented as SBOL-OWL terms or classes. For example, the `SequenceConstraint` term is used to restrict the relative ordering of subparts and the `MapsTo` term can connect hierarchical design information.

### 2.2 Encoding entities that are not used in SBOL serialization

SBOL-OWL includes additional terms that are not serialized as SBOL entities. Some of these entities are simply used to classify other SBOL entities and would be useful when querying genetic design information. However, such queries are currently not possible since the exact graph representation of data do not include any reference to these entities. For example, `TopLevel` is not a specific SBOL entity that would be present when designs are serialized. Unified Modeling Language (UML) (*26*) diagrams in the SBOL specification often utilizes such entities when describing entities with common properties. These interface classes are crucial for semantic reasoning and to simplify queries, and hence are included as classes in SBOL-OWL.

Other examples of such interface classes include `Identified`, which is defined to be any SBOL entity that can be identified uniquely using URIs. The use of interface classes in SBOL-OWL also allows identifying other entities through *is a* and *subclass* relationships. For example, `ComponentInstance` in SBOL describes how design elements can be defined using inputs and outputs to create hierarchical designs and modules. However, the `ComponentInstance` entity is not used in serialization. Instead, `FunctionalComponent` and `Component` entities in SBOL are derived from `ComponentInstance`, and are respectively used to serialize information at the structural DNA level to describe sequence composition of parts and at the functional level to describe molecular interactions. In SBOL-OWL these child entities are represented as classes that are subclasses of `ComponentInstance`, which can therefore be used in queries.

### 2.3 Representing relationships between design components

One of the reasons to develop SBOL-OWL is to validate SBOL documents using an ontological approach, where logical axioms are used to confirm that the ontology is consistent when it is merged with SBOL documents. This semantic layer provided by the ontology helps identifying inconsistencies through existing reasoners. To enable this approach, it is crucial to capture complex relationships between different SBOL entities ontologically. In addition to modeling the SBOL data model as an ontology, we developed SBOL-OWL to also capture complex validation rules in SBOL, as a set of logical axioms to prevent inconsistencies between the SBOL data model and genetic circuit designs.

Based on the serialization examples in the SBOL specification, relationships between SBOL entities are mainly modeled as *object* properties in SBOL-OWL. These object properties are defined with the `owl:domain` and `owl:range` properties respectively to indicate which SBOL entities would have these properties and which SBOL entities would these properties point to in order to link different SBOL entities. Some of the properties of SBOL entities can be literal values such as strings or numerical values. Such properties linking SBOL entities to literals are modeled as *data* properties in SBOL-OWL. However, if the value is a URI, such as an identifier to an external ontology term, the property is modeled as an object property too.

SBOL specification enforces strict rules to specify the cardinality of properties and whether they are required or not. The following rules were applied to represent these complex relationships between entities and values using SBOL-OWL properties, where possible. These rules can further be applied to extend SBOL-OWL consistently for future versions of SBOL.

- *Exactly one value*: The *domain* and *range* restrictions of the property are defined to capture the relationships between an entity and a value. The property is then defined to be *functional* to indicate that the entity can have at most one value. The class representation of the entity is then restricted to have at least one use of the property through the *some* (*someValuesFrom*) constraint. For example, a `Participation` entity in SBOL must have a `participant` property and the value must be an instance of `FunctionalComponent`. In SBOL-OWL, the `participant` property is defined to be functional so that there can be at most one `FunctionalComponent` for a `Participation` entity. Moreover, the `‘participant some FunctionalComponent’` restriction indicates that there must be at least one `FunctionalComponent` for a `Participation` entity. Finally, the `participation` property’s *domain* and *range* restrictions point to `Participation` and `FunctionalComponent` classes respectively.
- *Zero or one value*: Similar to the case above, the property is defined to be *functional*, and *domain* and *range* restrictions are defined to link entities and values. For example, the `version` property of the `Identity` is an optional `String` and can have at most one value. Therefore this property is defined to be a *functional* data type property. The *domain* and *range* restrictions point to the `Identity` class and `xsd:string` respectively.
- *zero or more value*: The property’s *domain* and *range* restrictions are defined to link entities and values. For example, a `ComponentDefinition` entity may have zero or more `SequenceConstraint` entities. Therefore, the `sequenceConstraint` object property’s *domain* and *range* restrictions point to `ComponentDefinition` and `SequenceConstraint` classes respectively.
- *one or more*: The *some* restriction on the entity is used to indicate at least one relationship. The property’s *domain* and *range* restrictions point to the entity and the value respectively. However the property is not defined to be *functional*. An example is the modeling of the relationship between `SequenceAnnotation`s and `Location`s. Each `SequenceAnnotation` can have at least one `Location` via the ‘`location some Location`’ restriction.

In addition to these rules, additional logical axioms are used to represent more complex validation rules where possible. These axioms are included as subclasses that restrict the definition of SBOL entities. For example, when describing hierarchical designs and connecting inputs and outputs of different design components, the component referred to as *“the remote”* must have the `access` type set to `public`. In SBOL-OWL, this constraint is split into logical axioms. Since the `remote` property is required, it is defined as *functional*. Using the existential *some* relationship, “`remote some ComponentInstance`“ axiom is defined. Finally, to restrict the potential values of the `access` property for `ComponentInstance`s, we use the clojure axiom `“remote only (access some public)”`.

To increase the flexibility of querying mechanisms using SBOL-OWL, we also defined inverse properties. For example, in addition to linking a `ComponentDefinition` and a `Component` entity using the `component` property, `isComponentOf` property is also defined, as an inverse property of the former. These inverse properties make the construction of logical axioms easier.

### 2.4 Incorporating external ontological terms

SBOL uses references to many other ontologies to provide the meaning of design entities where possible. Some of these external terms are also included in SBOL-OWL. For example, BioPAX terms `Complex`, `DnaRegion`, `RnaRegion`, `Protein` and `SmallMolecule` are used to indicate types of design components. In addition, Sequence Ontology (*27*) terms indicate the role of DNA-based components in designs. Similarly, EDAM (*28*) terms indicate types of external documents that are referred to and the Systems Biology Ontology (SBO) (*29*) terms are mainly used to classify biological interactions. The Provenance Ontology (PROV-O)(*30*) is also incorporated into the SBOL standard to capture information about design-build-test activities in order to track incremental changes in designs and the reasons affecting these changes.

In SBOL-OWL, SBOL entities are also defined as PROV-O entities. Therefore, provenance-based relationships can directly be used in semantic inferencing. In the SBOL specification, the base `Identity` class is used to describe common properties of SBOL entities and one of these properties is `provo:wasDerivedFrom`, which indicates how an SBOL entity is created. In the PROV-O, the `wasDerivedFrom` property is used between individuals of the `provo:Entity` class. Therefore, in SBOL-OWL, the `Identity` class is defined as a subclass of `provo:Entity`. This approach provides a seamless integration to integrate SBOL and PROV-O compliant tooling. Through ontological integration, other PROV-O related information such as activities, plans, and software agents can also be linked together.

### 2.5 The SBOL vocabulary

Not everything that is necessary to define genetic circuits exists in other ontologies. Therefore, the SBOL standard has been extended with specific terms where necessary. The SBOL specification describes these values as URI constants. For example, the `direction` and `access` properties are useful to respectively describe the direction of inputs and outputs, and whether they can be accessed by other designs. In SBOL-OWL, each URI is represented as a class to facilitate the execution of semantic queries. A parent class is also created for a set of related terms that are potential values for a specific SBOL property (Figure 3). For example, `private` and `public` are used as values of the `sbol:access` property when serializing SBOL documents. In SBOL-OWL, `public` and `private` classes are subclasses of the `Access` class. The `access` object property is created in such a way that only entities deriving from `ComponentInstance` can have this property and the *range* is strictly restricted to subclasses of the `Access` class (Figure 2). Similarly, we defined the `Direction`, `Orientation`, `Refinement`, `Restriction`, and `RoleIntegration` classes, each of which has subclasses that are used to describe genetic circuits.

**Figure 2:**
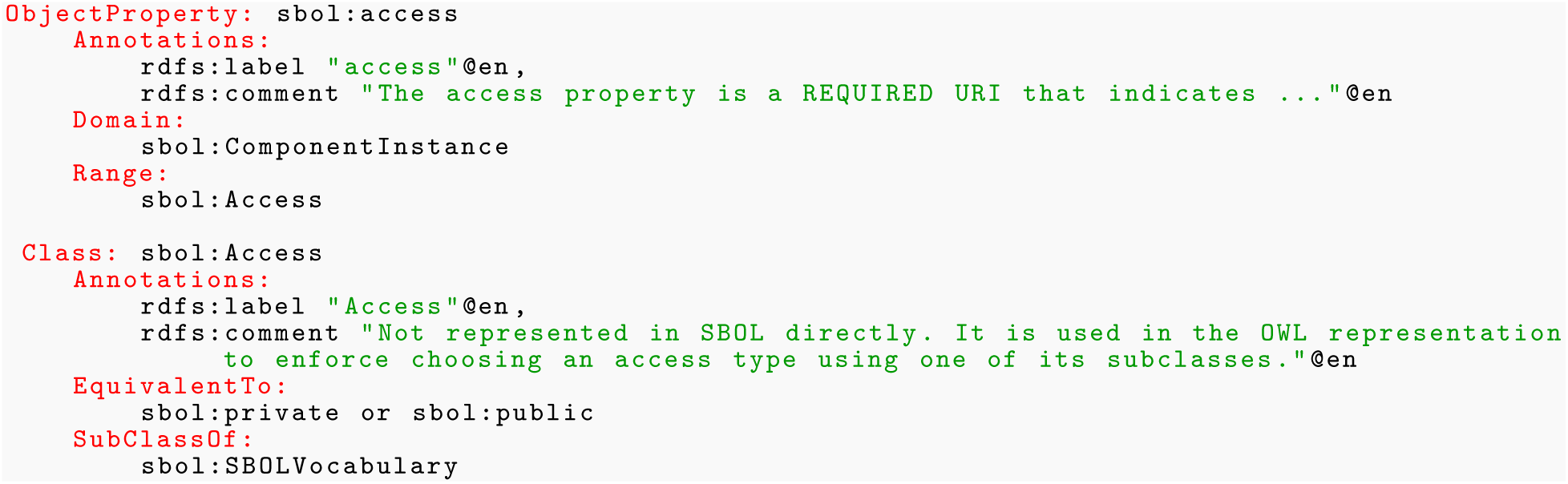
The definition of the `access` property and the `Access` class. Definitions are shown using the OWL Manchester syntax (*31*).

**Figure 3:**
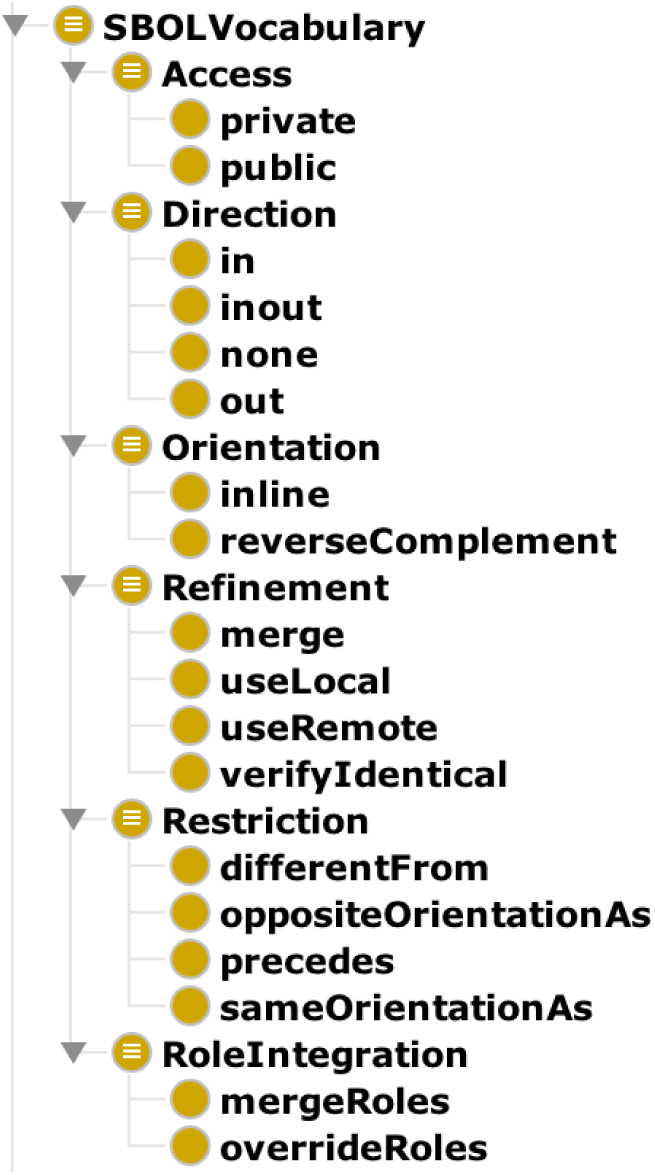
SBOL terms specific to the serialization of genetic circuit designs were described as OWL classes.

### 2.6 Adding metadata classes

SBOL is quite a verbose language. Although the data model is very flexible, it may require several statements to describe a biological concept. For example, a promoter term is represented using SBOL’s generic `ComponentDefinition` term. As a best practice, this entity has the value of `DnaRegion` from the BioPAX ontology for the `type` property, and the value of `SO_0000167` (promoter term) from the Sequence Ontology for the `role` property. The situation makes writing complex queries even more challenging when such entities are queried in the context of different information. The minimum information required to describe a promoter part can be simply represented as a single logical axiom, such that an entity is a “Promoter”. To facilitate the creation of such logical axioms, SBOL-OWL defines metadata classes (Figure 4A). For example, the `Promoter` metadata class would therefore indicate that any entity belonging to this class would have the `type` property set to `biopax:DnaRegion`, and the value set to `so:SO_0000167` (Figure 4B). Such metadata classes remove redundancies in representing data and enables writing simpler queries.

**Figure 4:**
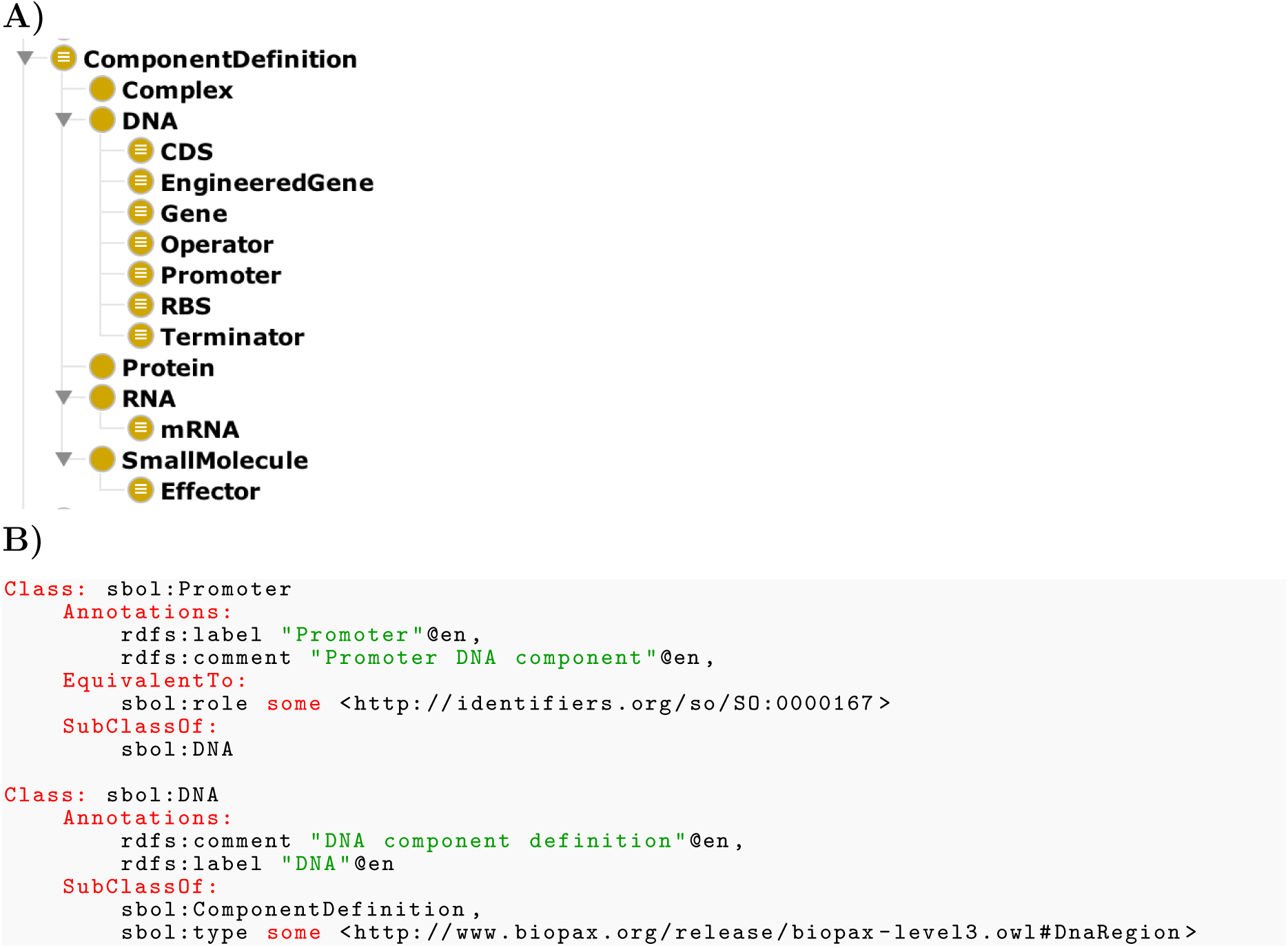
A. Metadata classes facilitate the creation of logical axioms. These classes often have human readable names and therefore queries are relatively easier to write compared to using identifiers from external ontologies and linking these identifiers to other SBOL specific terminology. B. Metadata classes include `Promoter` and `DNA`, which are shown using the OWL Manchester syntax. The `Promoter` class is defined to represent all entities that have the `sbol:role` property set to the `SO:0000167` promoter term from SO. It is a subclass of the `DNA` class that represent DNA entities. These entities are of type `ComponentDefinition` and have the `sbol:type` property is set to `biopax:DnaRegion`.

Metadata classes that are defined as subclasses of SBOL’s `ComponentDefinition` include `Complex`, `DNA`, `RNA`, `Protein` and `SmallMolecule`. These metadata classes correspond to Table entries in the SBOL specification. Each of these classes represent a different type of a design component which can have sequence information in a specified format. SBOL-OWL also enforces the types of sequences that can be associated with each type. Furthermore, we defined `CDS`, `EngineeredGene`, `Gene`, `Operator`, `Promoter`, `RBS` and `Terminator` classes as subclasses of the `DNA` class to implement SBOL’s best practices.

### 2.7 Querying in SBOL-OWL

SBOL data typically include information about design components that are considered as individuals in SBOL-OWL. These individuals belong to classes in the ontology. Querying in SBOL-OWL is therefore achieved through instance-level inferencing by using complex relationships and restrictions between these classes.

In SBOL-OWL, as in OWL more generally, one queries the data by creating a formal class definition, which then allows standard OWL inference engines (*22–24*) to determine membership into this class, thereby providing an answer to the query. For example, to find all instances of promoters, one would define the class `Promoter` exactly as in Figure 4B, and ask for all members (or instances) of this class. As described in the previous section, SBOLOWL includes a number of these common queries, built-in as metadata classes. In Figure 5C, we provide a slightly more complex query example, asking for all DNA components that have some child components.

**Figure 5:**
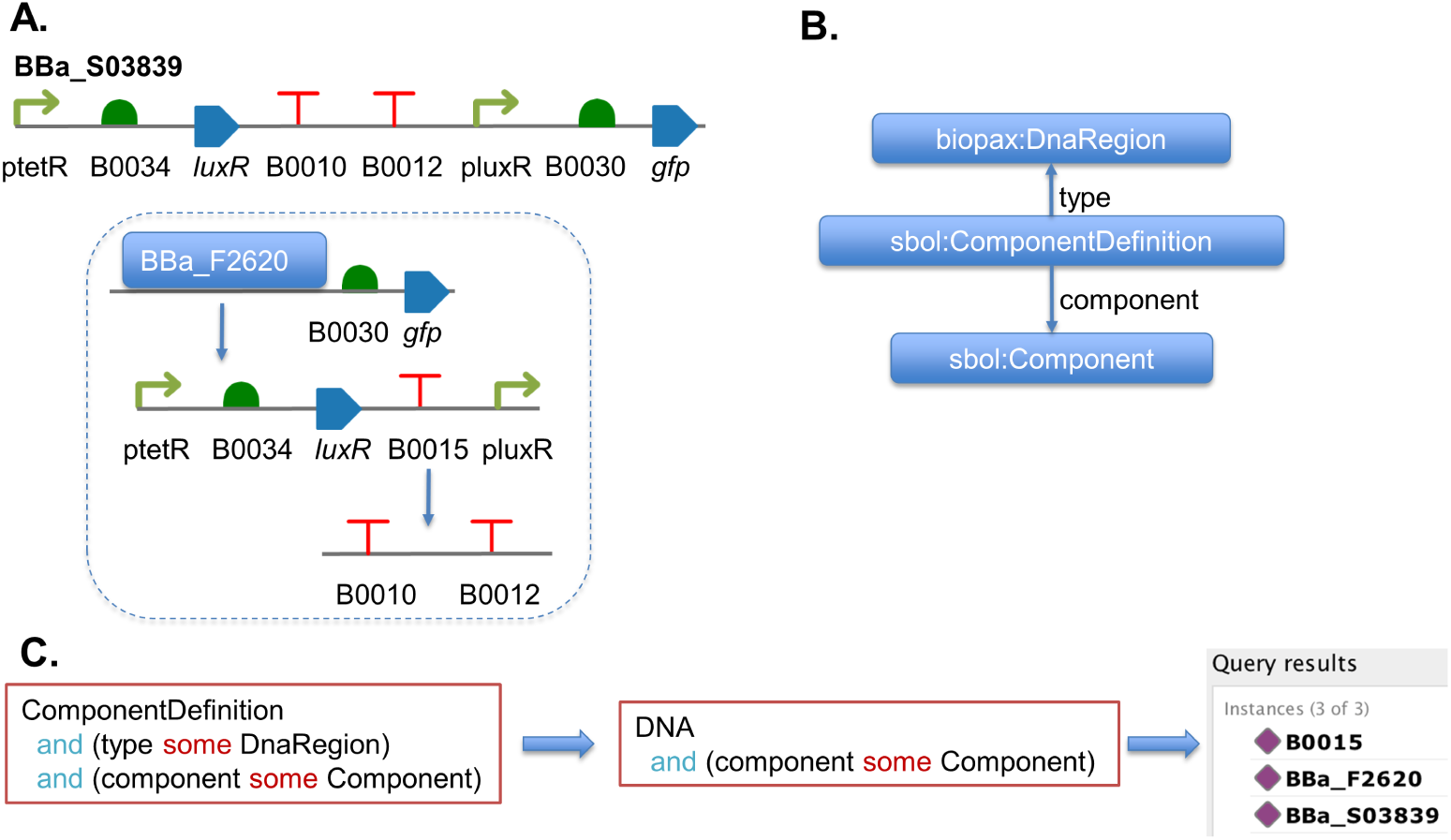
A. A genetic design example. B. SBOL data model to represent part-subpart relationships for DNA-based parts. Parts are represented using the `ComponenDefinition` entity. A `Component` entity stores the reference to the child part and is connected to parent part via the `component` property. C. SBOL-OWL based ontological queries.

As shown in Figure 5C, because queries are themselves classes, they can easily be reused in different queries. This is exactly the motivation by pre-creating metadata classes such as `DNA` and `Promotor` (Figure 4B). Moreover, a query class can refer to itself, thereby creating a recursive query, which are ideal for hierarchical designs such as are often used in SBOL. In contrast, SPARQL queries do not allow for recursion, and thus could only retrieve such information via multiple programmatic calls.

The challenge of querying SBOL hierarchical designs is one of our key motivations behind the design of SBOL-OWL. In SBOL, the relationship between a child and a parent part is not direct. Further, a part can be used in several different designs, and its use may have different properties. Thus, the class `ComponentDefinition` in SBOL refers to a part, whilst a `Component` refers to the use of that part by its parent (linked by the `component` property). These complex part-whole relationships become especially tedious to query as the size and complexity of designs increase. SBOL-OWL provides an ontological solution to query these complex designs. In the next section, we demonstrate how this is achieved.

## 3 Semantic reasoning for genetic circuit designs

The SBOL-OWL ontology is useful not only to formalize the standard representation of genetic circuit designs, but also to exploit the use of description logics to infer information about designs. Here, the ontology is used to achieve semantic reasoning. Typical competency questions to measure the validity of SBOL-OWL involve queries to extract information based on common properties of design entities. A simple example would be to retrieve all DNA parts which have subparts. A more formal description regarding this competency question is to return a list of `ComponentDefinition`s that of type `DnaRegion` and that have `some Component`s. Consider the design^3^ in Figure 5. The design consists of the BBa F2620 PoPS receiver device (*32*), followed by a RBS and a CDS encoding for the GFP reporter. The terminator, which is included in BBa F2620 consists of two individual terminator parts, and hence it is a double terminator. Therefore, the query should return the entities for the design, the PoPS receiver device and the double terminator.

Queries in SBOL-OWL typically utilize the *some* (*SomeValuesFrom*) restriction, since it is commonly used to capture relationships between SBOL entities. A *some* restriction indicates that there is at least one relationship and does not rule out other possibilities (*33*).

The OWL statement in Figure 5C represents SBOL entities that act as DNA-based composite parts, that are formed of other parts. This statement can also be used as a query to classify all SBOL entities conforming to this description. As a result, this query is used to retrieve part definitions representing DNA and having some parts. The query in Figure 5C is further simplified via the use of domain specific metadata classes and becomes “`DNA and (component some Component)`“.

The use of these metadata classes is especially important in more complex queries. These classes provide a way of semantic collapsing to create simpler queries. A competency question to retrieve parts which contain promoters can be formulated as follows: How do I return `ComponentDefinition`s that have `some Components`, which are `Promoter`s and are of `type DnaRegion`? This question can be captured in the form of a SPARQL query which is a graph pattern. An example SPARQL query is shown in Figure 6A. The query is constructed based on the SBOL data model, a subset of which is shown in Figure 6B. As it can be seen, SPARQL queries use exact graph relationships in order to report matching subgraphs in SBOL data. Even for this example, SPARQL queries can become quite complex using graph-based search mechanisms (Figure 6A-B). Such queries can be more easily constructed using SBOL-OWL (Figure 6C). Due to the way information is structured in SBOL-OWL queries are semantic. Additional `DNA` and `Promoter` classes are used to introduce domain knowledge without making changes in SBOL data. This semantic collapsing enables writing simpler and shorter queries via commonly used terms used in the synthetic biology domain.

**Figure 6:**
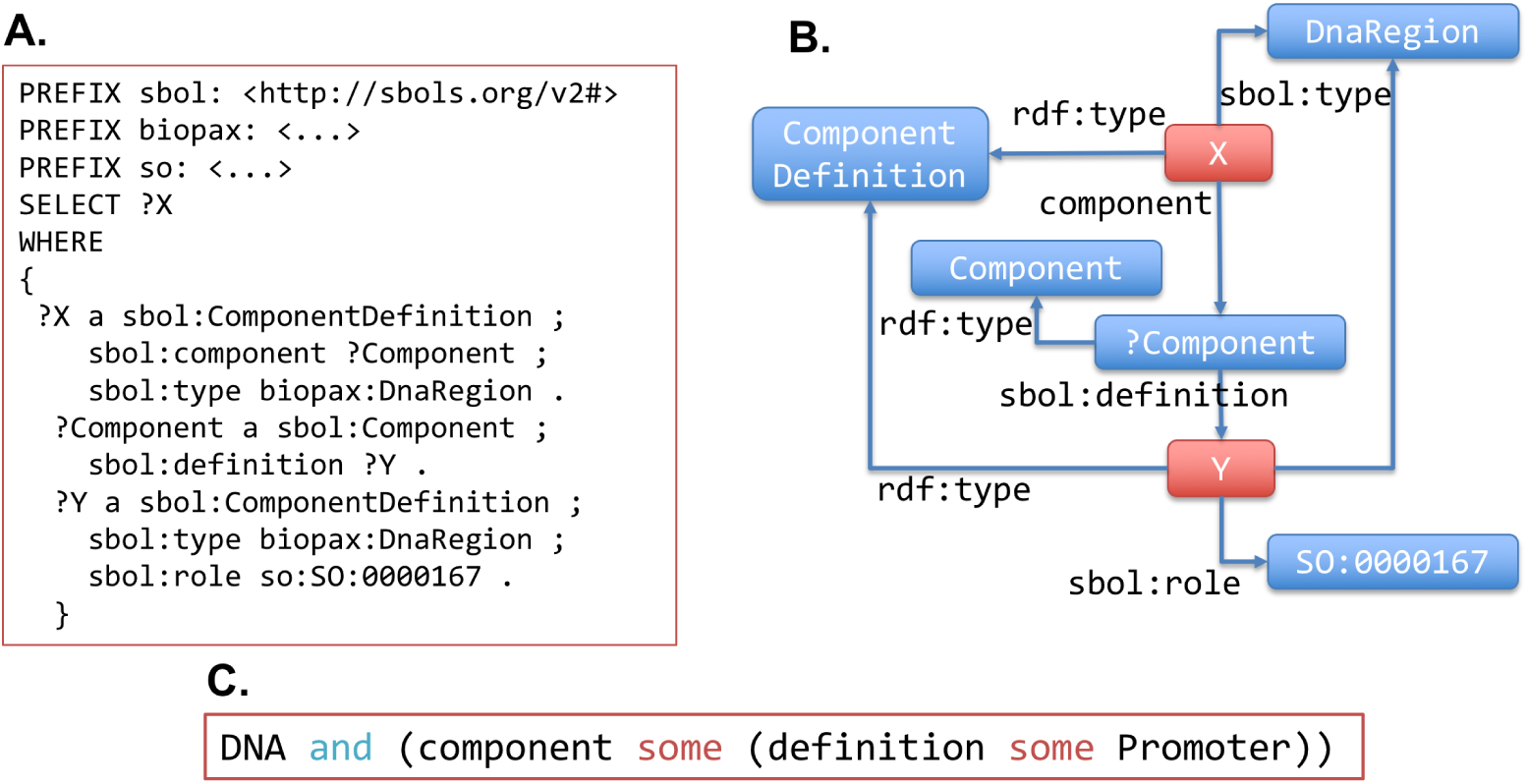
A. A SPARQL query to find parts that contain promoters. The query describes a graph pattern which is then used to search for matching design information. Parts and promoter subparts are represented with X and Y variables respectively. B. Graphical representation of the SPARQL query in A. C. The SBOL-OWL based representation of the SPARQL query shown in A. In this case, the query is created using the OWL syntax and logical axioms. Any `DNA` part that has `at least one component`, which is `defined` to be a `Promoter` is inferred from the design information. This query is captured as an OWL class which is then used by reasoners to classify parts. The result of the query is the list of individuals matching the class definition.

Considering the design in Figure 5A, neither the SPARQL nor the SBOL-OWL based query would list the BBa S03839 since this design does not include a promoter directly. However, the PoPS receiver device would be listed since it contains two promoters as first level parts. Clearly, this situation is not ideal and particularly a problem when using SPARQL queries.

SBOL-OWL terms and the concept of semantic collapsing can be used to create recursive queries to infer all of the required information. These queries can be constructed based on the transitivity of relationships using several classes and properties. This type of queries often involves querying based on a particular design component.

Recursive queries are especially needed when querying the ordering of biological parts. The location of a part can be crucial for the functioning of a genetic circuit. These locations affect *cis* interactions, which are about structural organization of DNA sequences. For example, for a genetic circuit to function as predicted, a promoter comes before a RBS to drive the expression of RNA molecules, and a RBS comes before a CDS to initiate translation. Positional constraints also affect the final concentrations and localizations of gene products and hence affect *trans* interactions. Therefore, it is important to gather information about a part with reference to other parts’ locations.

SBOL provides `SequenceConstraint` entities to represent relative ordering information between any two parts. For example, in order to represent the ordering of all parts in the PoPS receiver device, defining four `precedes` pairwise constraints is sufficient (Figure 7). Each constraint indicates which part precedes the other in the design.

**Figure 7:**
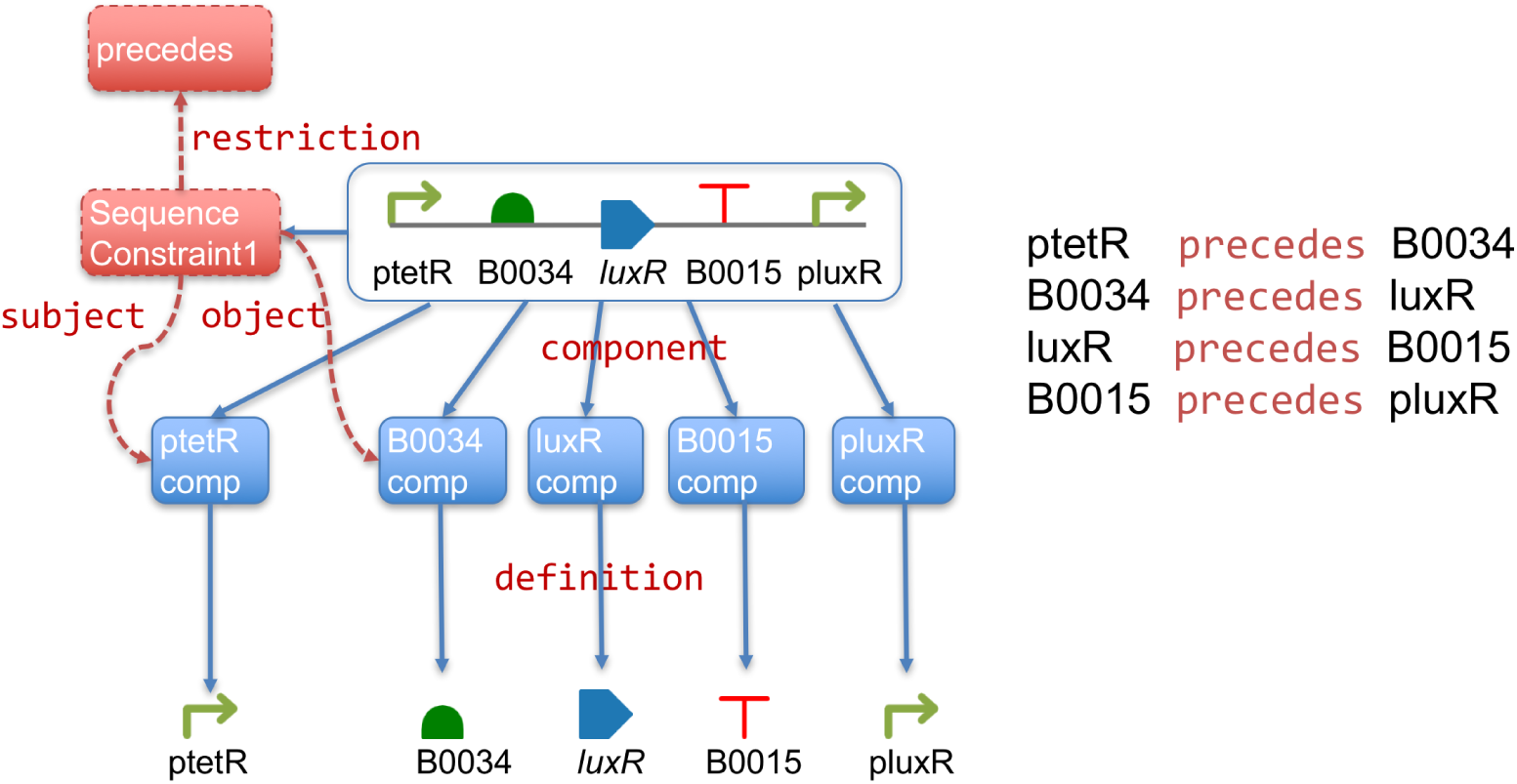
The PoPS receiver device consists of 5 parts. 4 pairwise precedes relationships is sufficient to represent the order of these parts. In the figure `sequenceConstraint1` is used to represent the relation between ptetR promoter and the B0034 RBS parts.

Finding the relationships based on the connection of nodes and edges using direct neighborhood is relatively easy and therefore parts proceeding others can be retrieved using SBOLOWL based queries. Figure 8A displays such a query to find the part that comes after the ptetR promoter. Here, we use the `value` (*hasValue*) (*8*) restriction in order to reference genetic parts in queries. However, it is not straightforward to find all parts that come after the ptetR part. Parts can be grouped semantically to indicate that any part coming after ptetR should be recursively queried. The latter approach is demonstrated in Figure 8C. The ptetRFollower is defined as an OWL class and its definition refers back to itself. When the class is submitted to reasoners, all parts that come after ptetR are considered. As a result, the query returns components for B0034, luxR, B0015 and pluxR.

**Figure 8:**
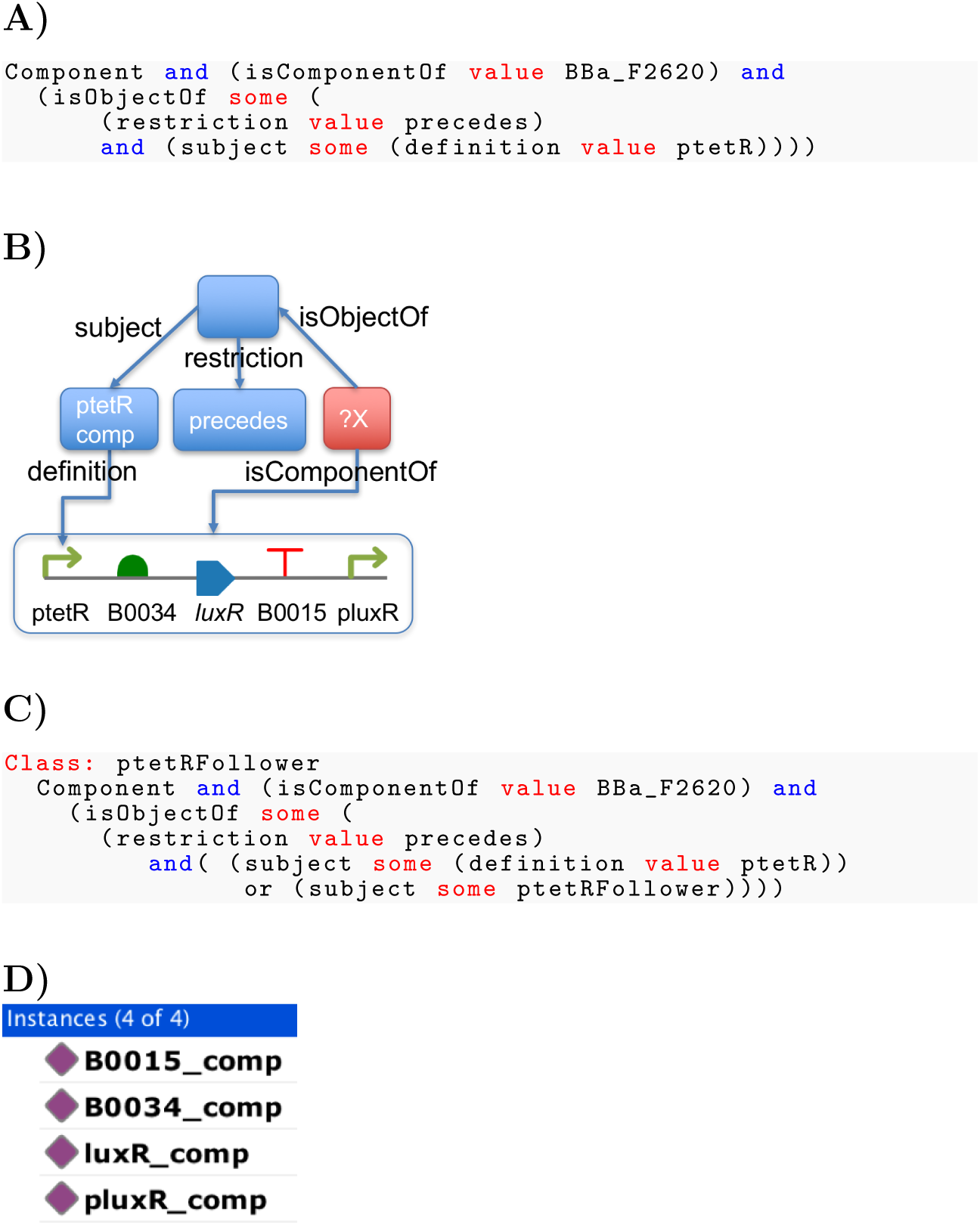
A. SBOL-OWL based query to find the part that comes after the ptetR part. B. Graphical representation of the same query. C. Recursive version of the query which is defined as a class to return all parts that come after ptetR. D. Results of the recursive query in C.

Even this approach may not be sufficient when parts are included in designs that include also other parts. Consider the design in Figure 5A. The query in Figure 8C would not be able to retrieve the B0030 RBS and gfp CDS parts, since they are not not included in the parent design of ptetR. The solution to this problem is to update SBOL-OWL based queries and to incorporate parent designs in queries. Such an approach is invaluable to query hierarchical designs and to find uses of a single part. An example of finding the use of a part in all designs is shown in Figure 9. The `ptetRParent` class in Figure 9A is used as a query to find all uses of the ptetR parents and their use recursively. The query in Figure 8C is then updated to also refer to parents of the ptetR part. As a result, the query in Figure 9B returns all parts that come after ptetR, but also any that follow its direct or indirect parents recursively. The query result is shown in Figure 9C.

**Figure 9:**
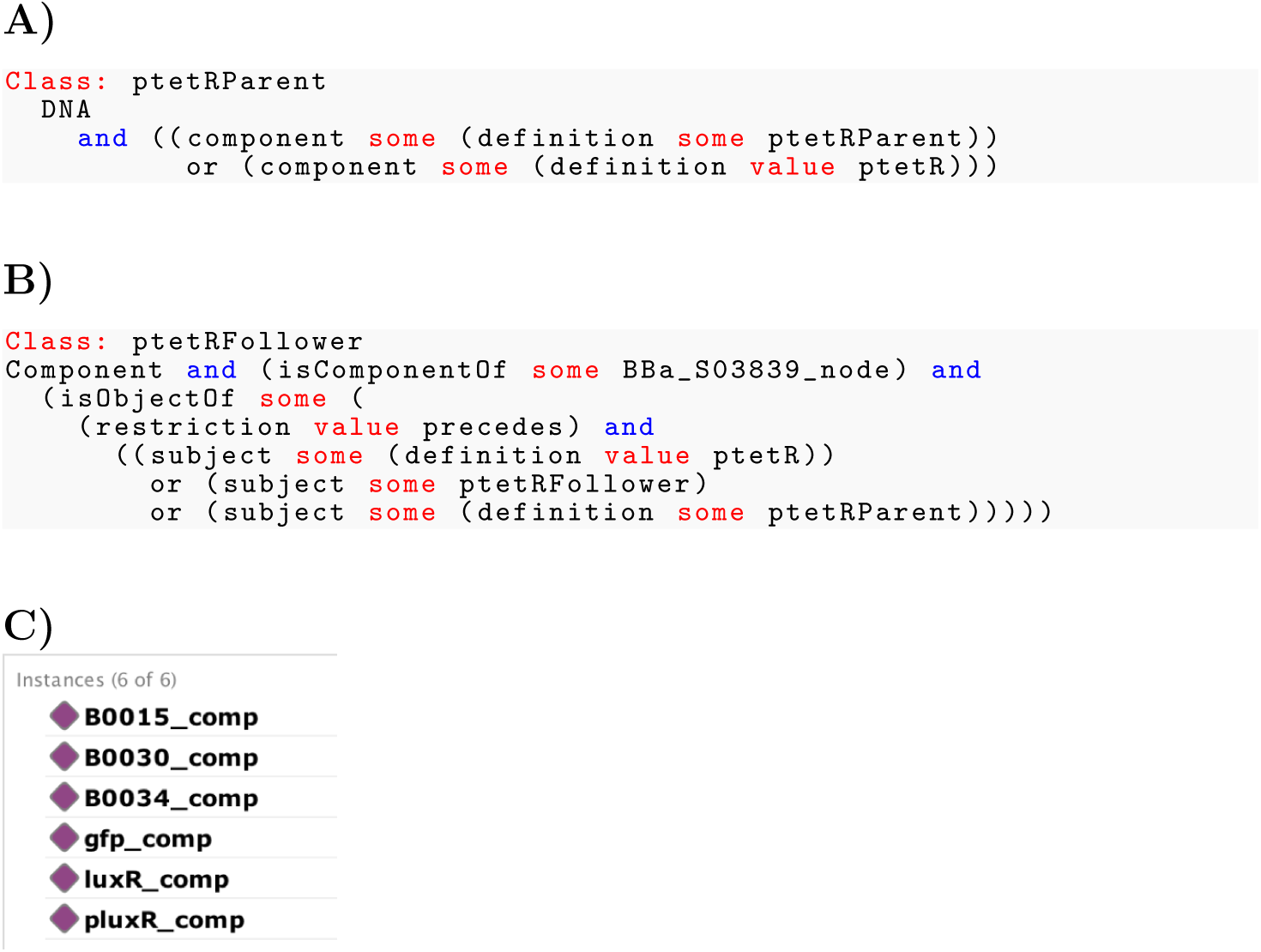
A. When ptetRFollower is submitted to reasoners, both BBa_F2620 and BBa_S03839 are inferred as designs where ptetR is used. B. The ptetRFollower class is updated to include designs where ptetR is used in order to recursively incorporate information about parent designs into the query. C. Results show parts that come after the PoPS receiver device (BBa_F2620) too.

## 4 Discussion

SBOL-OWL has been developed to complement the standardization efforts in synthetic biology. So far the community has focused upon a common syntax to facilitate the reproducibility of designs. Our goal here is to make information about genetic circuits machine understandable via an additional semantic layer. SBOL-OWL captures the SBOL specification as a Semantic Web ontology, standardizing both the syntax and semantics of the SBOL data model in a computationally tractable format. Semantic representation of genetic circuits has significant advantages and the potential to create new applications for synthetic biology. Examples include creating applications for data integration, visualization, model annotation and automated reasoning.

Here, we showed the utility of SBOL-OWL to infer information. The use of SBOL-OWL queries rather than complex patterns of graph-structure based SPARQL queries is more intuitive and efficient. Some of the inferencing tasks would be very difficult to implement using SPARQL. In some cases, multiple SPARQL queries would need to be executed, requiring the integration of results programmatically. On the other hand, multiple and complex graph queries which may be due to hierarchical representation of biological design information can be represented using relatively simpler ontological queries. Not all of the SBOL entities, such as `TopLevel` and `ComponentInstance` are included when genetic design information is serialized. As a result, these entities cannot be included in standard querying mechanisms. SBOL-OWL exploits the ontological representation of the SBOL specification and enables the use of any SBOL entity in queries. Additionally, SBOL-OWL introduces domain specific terms to simplify the construction of intuitive queries, removing redundancies in queries via parent-child relationships.

Representing genetic designs using RDF graphs already exploits existing Semantic Web tooling. Design information can be stored in RDF databases, called triplestores, and standard RDF libraries can be used to execute graph queries. SBOL-OWL adds a semantic layer over these queries. SynBioHub (*12, 13*) has previously successfully demonstrated the creation of instances of these triplestores. SBOL-OWL can potentially facilitate integration of data, distributed over multiple SynBioHub instances using a method called data federation (*34*). Using this approach, data are left in remote repositories, and a common semantics is used to integrate data from sub queries. Since SynBioHub is a graph database and stores SBOL data, SBOL-OWL can mediate the data integration process.

Semantic collapsing techniques can be exploited through metadata classes that are introduced in SBOL-OWL for commonly used design concepts. For example, a complex SBOL representation for a design component with the DNA type and a promoter role is simply represented via the Promoter class in SBOL-OWL. Moreover, hierarchical queries are simplified via class-subclass relationships. Child entities implicitly inherit properties of their parents. This approach can remove redundancies in queries and can also be adopted for data representation and visualization using SBOL in the future.

Semantic reasoning has huge potential to verify genetic circuit structures. Currently, constraints between any two DNA parts can be captured using SBOL. Although the terminology to define the relationship is not rich, efforts in this are are ongoing. SBOL-OWL can be easily integrated with the set of new terms in order to validate genetic circuits based on the order of DNA components.

As the SBOL specification is developed and new entities are added to cover even more types of biological data, it will be important to keep track of changes computationally and to create new libraries. Having the SBOL specification in the form of a machine accessible ontology will potentially facilitate the development of automated approaches to computationally verify and validate designs. SBOL-OWL already provides restrictions about how SBOL entities can be linked together and can be useful as a first step to verify genetic circuit designs. It can also be used to auto-generate libraries for tool developers.

SBOL-OWL can also facilitate utilizing large amount of biological data that can be mined for design information, through the integration of other ontologies. SBOL has already adopted the Provenance Ontology (PROVO)^4^ to provide provenance information about designs. SBOL-OWL exposes SBOL entities directly for PROV-O tooling. The SyBiOnt (*35*) has been developed as an application ontology and is promising as a way of providing an integration mechanism to different ontologies. The linking of SBOL-OWL and SyBiOnt can bridge the use of existing biological data and design information to create reliable biological applications. Since RDF is commonly used to capture semantic annotations for computational models in biology, SBOL-OWL will further facilitate the linking of these models and genetic circuit designs (*36–40*).

Currently, SBOL-OWL is limited to how the SBOL specification is developed. In the future, the specification can be tweaked slightly for more appropriate OWL formalization. This approach would simplify the serialization of SBOL into a bag of triples, whose semantics can be controlled by SBOL-OWL. The ontology is affected by some of the issues inherent in SBOL. Some of the properties such as “`role`“ are used to define relationships between more than two different entities and each representation may have different requirements.

SBOL-OWL is a useful resource for the community as a semantic layer for genetic circuit designs. Developers can use it to create new applications, for example to auto generate libraries, to provide easy-to-use tools for intuitively querying genetic circuits, to integrate data and so on. This ontology can play a key role to integrate huge amount of design information that is already available through the use of linked data and Semantic Web approaches.

## 5 Methods

### 5.1 Constructing the ontology

The SBOL-OWL ontology was programmatically constructed using Tawny-OWL (*41*), an API that provides high-level access to create entities for an ontology and to define the semantic relationships between those entities. Due to the built-in features of Tawny-OWL, SBOL-OWL has been developed using a textual user interface within an integrated development environment, which allows errors to be be identified when the resulting code is compiled. Moreover, SBOL-OWL utilises ontology patterns provided by Tawny-OWL. These patterns allows utilising ontology modelling solutions with less effort.

Clojure (*42*) was chosen as the programming language since Tawny-OWL is based on this language. Clojure is a functional programming language and Tawny-OWL exploits the advantage of this language to provide both a custom domain specific language and a set of macros to create ontologies. Here, additional macros were created in addition to using the Tawny-OWL API when creating the SBOL-OWL ontology. The resulting ontology was exported in OWL, RDF and Manchester Syntax (*31*) formats.

### 5.2 Testing the ontology with standard definition of genetic circuits designs

SBOL-OWL was specifically developed to provide a semantic layer for SBOL and hence we demonstrate how the ontology can be applied to genetic circuit designs that are represented using SBOL. SBOL-OWL does not require any changes to be made in SBOL files. However, a semantic reasoning process requires that information about genetic circuit designs and the ontological information defining semantic relationships between types of entities (or classes) are integrated and presented to reasoners together. To demonstrate this idea, we provide a Java application to merge the RDF graph content of an SBOL file and the RDF graph representation of the ontology. The resulting graph essentially stores classes from the ontology and individuals that come from the design, and can be submitted to existing reasoners directly. The Java application was developed using the Jena RDF library^5^ and libSBOLj (*43*) to process RDF graphs and SBOL data respectively.

### 5.3 Automated reasoning

Here, Protégé was used to execute semantic queries via built-in reasoners. However, the resources provided here are based on Semantic Web technologies and any tool supporting reasoning can be used. For example, Tawny-OWL can also be used programmatically to execute these queries in the form of a Clojure file. In this work, simple queries were executed directly as Description Logic queries via Protégé, and complex queries were defined as ontology classes.

### 5.4 Availability

The resulting ontology files and the source code to produce the ontology together with the Java library and examples demonstrating the application of the ontology for genetic circuit designs are available from http://sbolstandard.org/ontology.

## Acknowledgement

Authors thank Jacob Beal and Richard Markeloff for their valuable comments. The work of AG-M is supported by the SynBio3D project of the UK Engineering and Physical Sciences Research Council (EP/R019002/1) and the BioRoboost Contract of the European Union (820699). The work of CM is supported by the National Science Foundation under Grant No., 1522074, CCF-1218095, DBI-1356041, and DARPA FA8750-17-C-0229. Any opinions, findings, and conclusions or recommendations expressed in this material are those of the author(s) and do not necessarily reflect the views of the funding agencies.

## Graphical TOC Entry

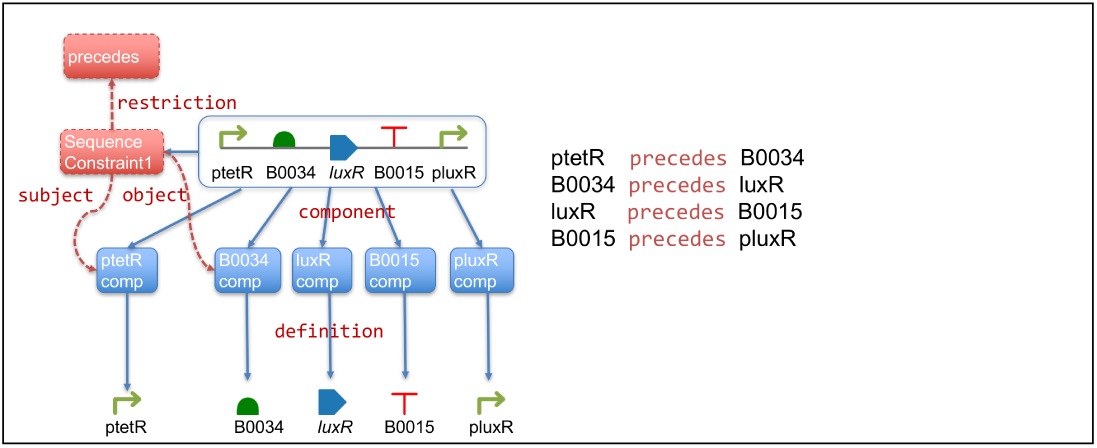

## Footnotes

^1^http://www.w3.org/TR/rdf-sparql-query

^2^http://www.w3.org/2004/OWL

^3^http://parts.igem.org

^4^https://www.w3.org/TR/prov-o/

^5^https://jena.apache.org

